# The Role of Semantic Inhibition in Aesthetic Appreciation: fNIRS Evidence from Poetry Reading

**DOI:** 10.1101/2024.10.08.617173

**Authors:** Huishu Liu, Xiaomeng Xu, Wanyan Sun, Dan Zhang, Yu Zhang

## Abstract

**Background:** Aesthetic education is pivotal in shaping a comprehensive and harmonious humanity. However, the transition from semantic comprehension to aesthetic appreciation remains poorly understood. This study, informed by transactional theory, sought to elucidate the cognitive mechanisms of aesthetic reading by examining its neural dynamics through functional Near-Infrared Spectroscopy.

**Methods:** Participants were tasked with reading Tang poetry aesthetically, with fNIRS monitoring brain activity in frontal and temporal regions.

**Results:** Compared to an efferent reading task, the aesthetic reading task revealed a distinct three-stage neural dynamic pattern. Initially, aesthetic reading showed similar HbO activation in all regions, likely indicating the semantic processing phase. This was followed by a divergence, with a decrease in HbO over the left primary somatosensory cortex and the left superior, inferior, and middle temporal gyri, suggesting inhibition of semantic processing. Finally, a resurgence of activity in these areas was observed, along with an increase in HbO over the left dorsolateral prefrontal cortex, which could be associated with memory, imagination, and empathy. This reactivation correlated with participants’ self-reported aesthetic appreciation scores.

**Contribution:** The findings reveal the temporal and spatial dynamics of brain activity during aesthetic reading, enhancing our comprehension of underlying cognitive processes.

## 1 Introduction

Aesthetic education is pivotal in shaping a comprehensive and harmonious humanity, especially in modern society (Schiller, 2016). It brings into harmonious operation the various faculties of human beings—emotion, reason, imagination, sensibility, moral judgment (Csikszentmihalyi, 1997; Dewey, 1934; Gardner, 2011; Goleman, 1995; Greene, 2000), and social engagement (Vygotsky, 1978) and cultural awareness (Nussbaum, 1997; Ştefan-Sebastian, 2014). Aesthetic appreciation takes various forms, such as appreciating paintings, listening to music, or reading. This study focuses specifically on the aesthetic appreciation of poetry, particularly reading classical Chinese poems. Reading plays a key role in both formal education and informal learning environments. The aesthetic appreciation of poetry is not only accessible to most students but also a fundamental component of reading classes and liberal arts education. It fosters cognitive skills like critical thinking, as well as non-cognitive attributes such as imagination and empathy (Catterall, 1998; Greene, 1978).

Grounded in the transactional theory (Rosenblatt, 1994), an influential framework in aesthetic reading in educational practice, this study employs neuroscience-based methods to explore the cognitive mechanisms underlying aesthetic reading. The transactional theory distinguishes reading behaviors into efferent reading and aesthetic reading through the concept of “reader’s stance”. It suggests that the choice of reading approach depends on the reader’s stance, determining whether the reader focuses on “efferent” elements such as vocabulary meanings, plot content, and objective information, or on “aesthetic” elements emphasizing personal emotional experiences and imaginative associations (Rosenblatt, 1986). Compared to efferent reading, aesthetic reading places significant emphasis on the fusion of personal experiences with textual fragments, thereby forming a personalized connection between the reader and the text (Robinson, 2020). Through imagination, association, and anticipation, readers construct meaningful interpretations of the text, leading to subjective assessments or evaluations (Fowles et al., 1977). Readers in the stance of aesthetic reading are more inclined to engage in an immersive reading experience (Graves, 2011), which also triggers intense emotional fluctuations and resonant responses (McEneaney et al., 2009). Extensive empirical evidences indicate that readers who engage in aesthetic reading develop a deeper and more nuanced understanding of texts and are more inclined to make positive evaluations (Blakemore, 2012; Pilonieta & Hancock, 2012)

However, despite the theoretical analysis of what aesthetic reading should be, existing theories did not explain how to achieve it from the perspective of an inner, cognitive and emotional process. In aesthetic reading, according to transactional theory, external reference and internal response are two procedural conditions for generating meaning in aesthetic reading. External reference involves the reader processing the text through the context of poetry and their existing linguistic symbol experience, identifying objects in the external world to understand the text’s meaning; internal response refers to the reader’s focus on language symbols in their inner world, accompanied by processes such as association, imagination, and empathy (Rosenblatt, 1994). Wolf also posits that readers need to first understand the basic meaning of the text, and then utilize their imagination and analytical skills to deeply understand the text (Wolf, 2018). Therefore, aesthetic reading is conceptualized as a two-stage process: the first involves comprehending external elements, such as language and semantics, while the second entails the deeper internalization of these elements, transforming them into an aesthetic experience (Rosenblatt, 1994; Rosenblatt, 2018). Understanding the trigger that shifts semantic comprehension into aesthetic appreciation is critical. Existing theories predominantly emphasize external factors influencing this process, such as enriching life experiences to enhance empathy or fostering critical, egalitarian thinking. However, they often overlook the underlying cognitive mechanisms driving this transition.

Besides theoretical discussions, empirical research on the educational application of the transactional theory basically demonstrated the value and teaching methods of aesthetic reading (Blakemore, 2012; Forouzani, 2017), rarely investigating the intrinsic mechanisms of aesthetic reading. In fact, it is indeed challenging to measure or self-report the subtle, intrinsic process of aesthetic reading by traditional methods.

To address this issue, neuroscience-based measurement methods may offer novel evidence and insights, as existing studies have provided cumulative knowledge that are helpful to link the two disciplines. From the perspective of cognitive neuroscience, mechanisms of aesthetic reading and efferent reading are distinctly different. Efferent reading lacks personal subjective characteristics and instead focuses more on objective information such as textual meaning, creative techniques, and linguistic details. Thus, efferent reading may involve cognitive processes such as language comprehension, text information processing, and semantic control. When individuals utilize semantic understanding and judgment to process textual information, areas such as the left inferior temporal gyrus, left superior temporal gyrus, and middle temporal gyrus are notably activated (Davey et al., 2016; Friederici et al., 2003; C. Whitney et al., 2011).

In contrast, the psychological processes involved in aesthetic reading are more focused on remembering, imagining, empathizing. Specifically, imagination activity is involved when the reader reconstructs stored memories into new structural images and engages in the iterative processing and contemplation under the stimulation of textual cues. Such imagination enables both cognitive empathy and emotional empathy for aesthetic appreciation (Brinck, 2018).

According to existing neurophysiological studies, the above-mentioned cognitive processes have been reported to be associated with certain brain regions. Autobiographical memory, which involves an individual’s recollection of their own experiences, is closely associated with the lateral and medial prefrontal cortex. The dorsolateral prefrontal cortex is also involved in the retrieval and encoding of episodic memories, showing hemispheric specialization. The left dorsolateral prefrontal cortex is more sensitive to encoding and retrieving linguistic information, while the right dorsolateral prefrontal cortex is primarily responsible for encoding and retrieving non-linguistic materials (Balconi, 2013). The activation of re-creative imagination primarily occurs in the left hemisphere of the brain. Among ordinary readers without literary training or innate literary talents, there is a higher level of activation observed in areas such as the left anterior cingulate gyrus in the left frontal lobe (Zhao, 2018). Cognitive empathy is primarily associated with the activation of the ventromedial prefrontal cortex, while emotional empathy is mainly associated with activation in areas such as the anterior cingulate cortex, anterior insula, anterior midcingulate cortex, amygdala, and secondary somatosensory cortex (Dodell-Feder et al., 2011; Mclatchie et al., 2016). The evaluation of textual aesthetics is closely linked to brain activity, with the activation level of the frontal cortex showing a significant positive correlation with aesthetic assessment (Jacobs, 2015).

From a temporal perspective, which is closely related to the mechanism of transcending from semantic understanding to aesthetic appreciation, the evidence is rare. Although different phases were proposed by neurophysiological studies, these models did not address the issue of transcendence. An “aesthetic trajectory” was proposed with three phases of familiarity, novelty, and synthesis (Fitch et al., 2018), but such phases are not in accordance with existing theories. Other studies discussed some issues related to phases, but their focus is not on the transcendence issue either (Höfel & Jacobsen, 2007; Leder, 2013). In order to further understand the dynamic processes underlying the transition from semantic understanding to aesthetic appreciation, neuroscience methods can provide valuable insights. Current research has identified the left dorsolateral prefrontal cortex as potentially being associated with imagination and aesthetic evaluation (Brothers, 1990; Light, 2009; Bernhardt, 2012; McNorgan, 2012; Engen, 2013; Yang, 2018), while the left primary somatosensory cortex, left superior temporal gyrus, left middle temporal gyrus, and left inferior temporal gyrus are likely involved in word processing, language comprehension, semantic knowledge storage, and semantic control (Aryani, 2018; Binder, 2009; Booth, 2001; Booth, 2006; Gaillard, 2001; Joseph, 2001; Whitney, 2011). However, no studies have yet explored the temporal dynamics involved in the transition from semantic understanding to aesthetic appreciation.

This study thus attempts to explore the transcendence mechanism from sematic understanding to aesthetic appreciation, through the lens of neural activities. It uses portable functional near-infrared spectroscopy (fNIRS) devices to analyze and compare brain activity differences between aesthetic reading and efferent reading, along the temporal dimension. fNIRS, with its high temporal resolution, moderate spatial resolution, and portability, is well-suited for capturing the dynamic neural processes of aesthetic reading, particularly in identifying the activation of relevant brain regions, making it an ideal tool for this study. In summary, this research aims to answer the following questions:

## 2 Materials and Method

### 2.1 Participants

A total of 35 university students were recruited (18 males and 17 females with an average age of 22.08 years). Participants were intentionally selected from diverse academic backgrounds, ensuring a balanced representation of disciplines including sciences, engineering, and humanities. All of the participants were exclusively right-handed and possessed either normal vision or vision that had been corrected to normal standards.

Device Setup and Calibration: During this phase, the experimenter assisted the participants in fitting the fNIRS equipment (NirSmart, Danyang HuiChuang Medical Equipment Co., Ltd., China), which included the fNIRS cap, emitter probes, and receiver probes. The experimenter calibrated the device to meet the experimental requirements. The device’s sampling rate was 11 Hz.

### 2.2 Stimuli

This study used poetry from China’s Tang Dynasty as the stimulus material, which possesses profound aesthetic value and serves as a type of literature that easily evokes readers’ imaginative and emotional resonance. Due to the rich variety of poetic forms in the Tang Dynasty, and to control for the influence of unexpected factors on research outcomes, this study focused exclusively on the Tang Dynasty’s five-character regulated verse (*wu yan lv shi* in Chinese pinyin). In the process of selecting experimental materials, this study not only considered the thematic diversity of the poetry, such as landscapes, farewells, frontier garrisons, historical musings, and travel, but also evaluated the poetry text materials from six aspects: reading difficulty, familiarity, emotional valence, arousal level, engagement, and aesthetic response degree (Gao, 2017). This was done to ensure a set of suitable and unbiased material.

Based on the above criteria, this study initially selected 25 poems sourced from the college students’ textbook “*A History of Chinese Literature*” (3^rd^ edition, volume 2)(Yuan, 2017). Then, to exclude any unsuitable poem, 13 graduate students majoring in education were invited to rate the 25 chosen poems. Ratings were conducted on a scale of 1 to 7 for all dimensions, including difficulty, familiarity, emotional resonance, level of engagement, aesthetic response, and imaginative difficulty. To check the consistent evaluation criteria among the 13 raters, Kendall’s coefficient of concordance reliability analysis was performed using SPSS 23.0. The Kendall’s coefficient of concordance reached a significant level (*p* < 0.001), indicating a high level of agreement among the 13 raters in their poem assessments. Based on the aesthetic response dimension, the top 20 poems with relatively higher levels of aesthetic response were selected as a refined set of materials. Experts in aesthetics were invited to double check if these 20 poems were appropriate for the experiment material. Overall, the 20 selected poems exhibited moderate levels of familiarity, emotional valence, engagement, and aesthetic response, and low levels of reading and imaginative difficulty. See Appendix A Table A1 for details.

### 2.3 Behavioral Assessments

A “Reading Stance Scale” was developed for participants to self-assess their aesthetic and efferent reading stances after the reading tasks. This scale was drew primarily from Mohammad’s aesthetic reading self-assessment scale (Forouzani, 2017) and was revised based on the essence of aesthetic and efferent reading, dividing it into two dimensions: aesthetic reading and efferent reading, comprising a total of 17 items. The scale uses a seven-point rating system. To test the reliability and validity of the scale, this study recruited a total of 38 university students in a pilot study. The participants were asked to adopt either an aesthetic or an efferent reading stance to read 10 Tang Dynasty five-character regulated verses. After the reading task, they were invited to fill out the “Reading Stance Scale” to report their degrees of aesthetic and efferent reading during that task. According to the principal component analysis, the 17 items of the Reading Stance Scale were extracted into exactly the two dimensions: aesthetic reading stance and efferent reading stance. The Cronbach’s alpha coefficients for the two dimensions were 0.92 and 0.78, respectively, both greater than 0.70. Therefore, this scale demonstrated good structure validity and internal consistency. See Appendix B Table B1 for “Reading Stance Scale”.

### 2.4 Protocol

The experiment (see Figure 1A for details) consisted of two phases: the preparation phase and the experimental phase. The preparation phase equipped the participants with the theoretical background necessary for the subsequent formal experiment. During the preparation phase, the experimenter informed the participants about the reading requirements and imparted foundational knowledge of the transactional theory, aiding the participants in understanding the concepts, characteristics, and application methods of aesthetic reading and efferent reading. As illustrated in Figure 1C, participants were required to wear fNIRS devices in the experimental phase. The procedure was divided into four parts: device setup and calibration, presentation of example tasks, main experiment, self-assessment questionnaire, and personal information report. The details are explained as follows.

**Figure 1.**
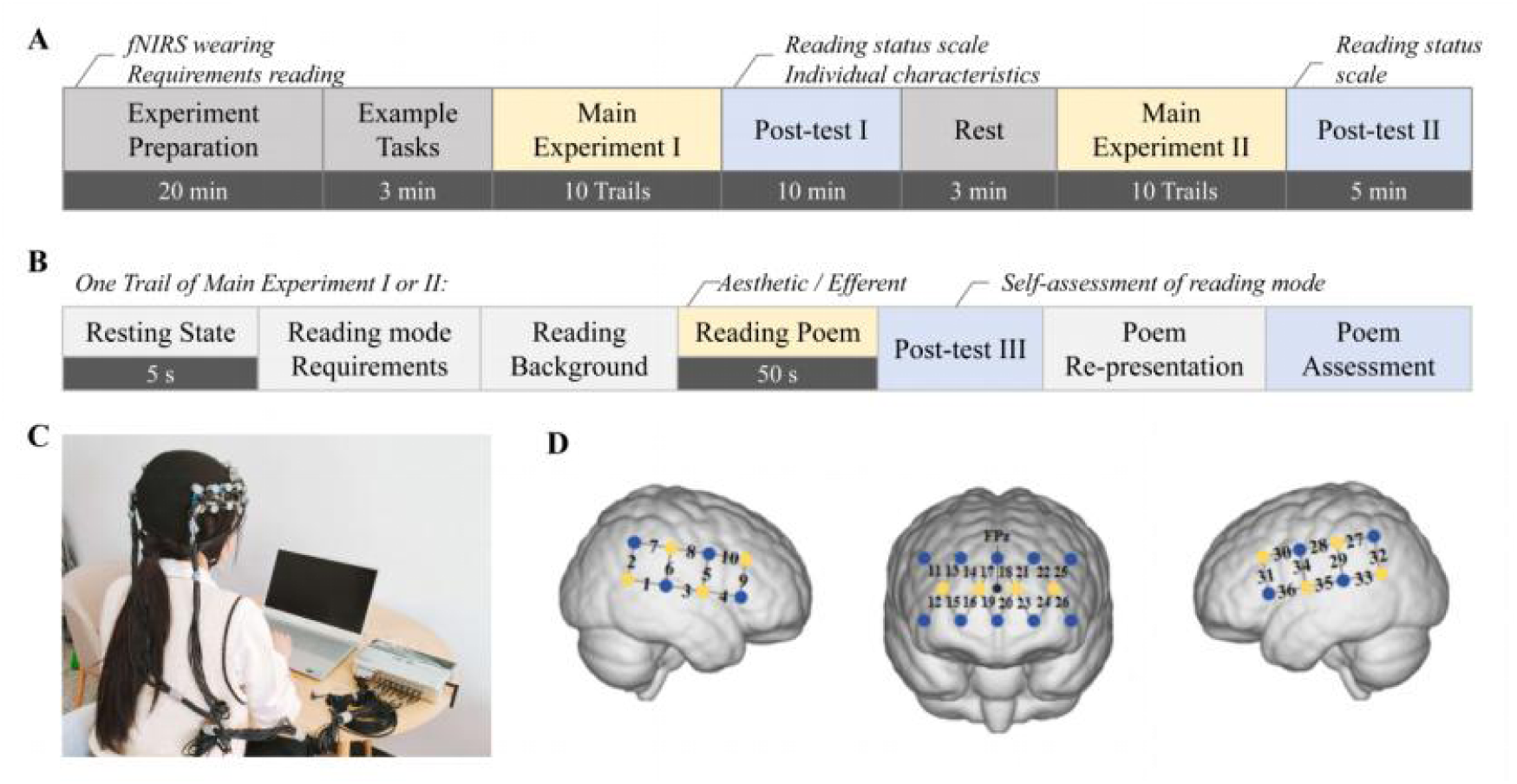
Experimental Design. **A,** the protocol of the formal experiment. **B,** one trail of main experiment round I or round II which was repeated for 10 times with the same reading stance in one experiment round. 20 poems were offered to participants randomly. **C,** a photo of the participant wearing fNIRS device in the experiment. **D,** fNIRS probe setup. The numbers represent the measurement channels.

#### Presentation of Example Tasks

In this phase, the experimenter ran the E-Prime software program on the computer. This program presented tasks corresponding to both aesthetic reading and efferent reading conditions. The experimenter explained the task requirements and the usage of the program for each reading condition to the participants. The participants practiced using the program and became proficient in responding to the tasks through the example tasks, which followed the same procedure as the formal experiment.

#### *The main experiment* was divided into two rounds

Round I and Round II. In each round of the main experiment, participants were required to consecutively complete the reading tasks of 10 poems using one of the two reading stances. The reading stances required in Rounds I and II were different. The order of reading stances (“Aesthetic-Efferent” or “Efferent-Aesthetic”) for the two rounds were randomly assigned for each participant. Each order accounted 50% across all participants’ tasks. Within each round, the presentation sequence of each poem was also randomly generated. The specific procedure is illustrated in Figure 1B.

The tasks for each poem included reading the poem, self-evaluating their reading stances, and evaluating the poem in terms of familiarity and likability. For each poem task, the program first presented either the “Aesthetic Reading Task Instruction” interface or the “Efferent Reading Task Instruction” interface based on the randomly generated order. Participants could proceed to the next interface by pressing the Enter key after reading the instructions. The program then displayed the “Poem Background Interface,” and participants could advance by pressing the Enter key. Subsequently, the program showed the “Read Poem Interface,” which included the original poem text and annotations for difficult-to-understand phrases.

Participants were required to read the poem using either aesthetic reading or efferent reading approach according to the task instructions. This 50-second poem reading segment constituted a pivotal timeframe in the subsequent analysis of fNIRS data. By examining the changes in the level of oxygenated hemoglobin (HbO) within this time window, we aimed to elucidate the distinct neural activities engaged by aesthetic and efferent reading.

After a 50-second reading period, participants were required to answer a series of multiple-choice questions. They were then asked to report the extent to which they adopted the instructed reading stance. If, during the reading of this poem, the participant reported using a reading stance opposite to the one instructed in the experimental protocol, the data pertaining to this poem were deemed inadmissible and accordingly excluded from analysis. Finally, the “Evaluate Poem Interface” prompted participants to rate the familiarity and likability of the poem on a scale of 1 to 7, where 1 represented “Very Unfamiliar/Disliked” and 7 represented “Very Familiar/Liked.” After the survey, the program presented a blank cross interface for 5 seconds as a resting-stage before moving on to the reading task of the next poem.

In order to understand the experimental conditions of the participants, as well as the overall reading stances during both rounds of main experiments, this study required participants to provide self-assessments of their overall reading stances using the Reading Stance Scale after each round (i.e., Round I and Round II).

### 2.5 fNIRS Data Acquisition and Processing

This study employed a 36-channel fNIRS system (NirSmart, Danyang HuiChuang Medical Equipment Co., Ltd., China) to measure participants’ neural signals during reading activities. As shown in Figure 1D, this cap layout encompassed brain regions intimately associated with language processing and reading comprehension, specifically targeting the frontal cortex and temporal lobe as key regions of interest (ROIs) based on previous studies. The probes formed a network structure, consisting of 18 sources and 12 detectors, totaling 36 channels symmetrically distributed across the left and right hemispheres. The center of the frontal cortex probe group was located at the midline of the frontal region (FPz), and according to the international 10/20 system, CH35 and CH3 were anchored at T3 and T4. The fNIRS signals were recorded at a sampling rate of 11 Hz, using near-infrared light wavelengths of 730 nm and 850 nm.

The NirSpark software (Danyang HuiChuang Medical Equipment Co., Ltd., China) was employed for preprocessing fNIRS signals, a tool that has been validated in previous studies (Ge et al., 2021; Tan et al., 2021). The preprocessing steps were as follows: (1) Irrelevant time segments were excluded based on markers, and only data from the experimental period were selected for analysis. (2) Based on participants’ feedback, fNIRS data from the time slots during which participants reported adopting a reading stance opposite to what was required were excluded. (3) A spline interpolation algorithm was employed to correct motion artifacts in the acquired signals across various channels. This widely adopted correction technique is advantageous in its ability to specifically target and amend pre-identified artifacts. (4) Motion artifacts were automatically identified and corrected. Signal changes exceeding 6 standard deviations from the entire time series were classified as motion artifacts. The amplitude threshold (AMP thresh) was set to 0.5 for automatic detection and correction. Any signal with an amplitude change less than or equal to 0.5 was deemed insignificant, noise-induced, or part of background fluctuations and was subsequently disregarded. The light intensity signals were then converted into optical density signals. (5) A band-pass filter (0.01 Hz - 0.2 Hz) was applied to remove physiological noise, including respiration and heartbeat. (6) HbO data were calculated using the modified Beer-Lambert law, converting optical density signals into HbO concentrations. This enabled the acquisition of HbO measurements at a sampling rate of 11 Hz across 36 channels for each participant during the 50-second reading period for each poem.

### 2.6 Calculation of the Temporal Dynamics of Brain Activity During Aesthetic Reading and Efferent Reading

To calculate the temporal dynamics of HBO levels in different brain regions during aesthetic and efferent reading, a paired t-test analysis at each time point was conducted. Given the intricate nature of the cognitive processes engaged during poetry reading, this study initially segmented the 50-second reading period into 1-second segments for each participant. The mean HBO levels per second for each poem were calculated for each participant. Subsequently, the HbO values obtained from the ten ancient poems were aggregated to derive a comprehensive measure of task-related HbO per second. Each participant provided data for 36 channels across 100 data points, with 36 * 50 data points corresponding to aesthetic reading and 36 * 50 data points corresponding to efferent reading. For each second of reading tasks, the activation level of HbO was computed as the difference between the task-stage HbO and the corresponding resting-stage average HbO. For instance, the activation level of HbO for aesthetic reading tasks was calculated as following:

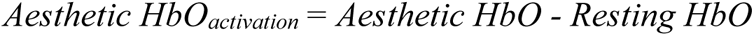

where *Aesthetic HbO* represents the HbO for aesthetic reading task and *Resting HbO* denotes average resting-stage HbO for aesthetic reading. Finally, a paired t-test analysis was performed to compare the activation levels of HbO between aesthetic and efferent reading tasks at each time point, thereby identifying the dynamic differences in brain region activation under the two reading stances. To ensure stability in the results, and considering that shorter durations might lead to unstable outcomes, this study focused only on results with a significant duration of 3 seconds or more.

## 3 Result

### 3.1 Behavioral Task Performance

According to Figure 2A, there are significant differences in participants’ degrees of aesthetic reading stances and efferent reading stances under the two experimental reading conditions (*p* < 0.001). Firstly, under the requirements of the aesthetic reading experiment, participants’ degrees of aesthetic reading stances (M= 5.34 ± 0.67) were significantly higher than those under the requirements of the efferent reading experiment (M= 3.02 ± 0.99). This suggests that the aesthetic reading experiment requirements effectively activated participants’ aesthetic reading stances. Secondly, under the requirements of the efferent reading experiment, participants’ degrees of efferent reading stances (M= 5.64 ± 0.53) were significantly higher than those under the requirements of the aesthetic reading experiment (M = 3.47 ± 0.96). This indicates that the efferent reading experiment requirements effectively activated participants’ efferent reading stances. In addition, there is no significant difference in the accuracy of participants’ responses between aesthetic reading tasks (M= 6.29 ± 1.69) and efferent reading tasks (M= 6.40 ± 1.48) (*p*=0.76). There was no significant difference in familiarity levels between any two pairs of the four poem sets (F_(3,66)_=0.09, *p*>0.05) according to one-way analysis of variance (ANOVA). The detailed information and the participants’ average level of engagement are listed in Appendix A Table A2, A3 and Table A4.

**Figure 2.**
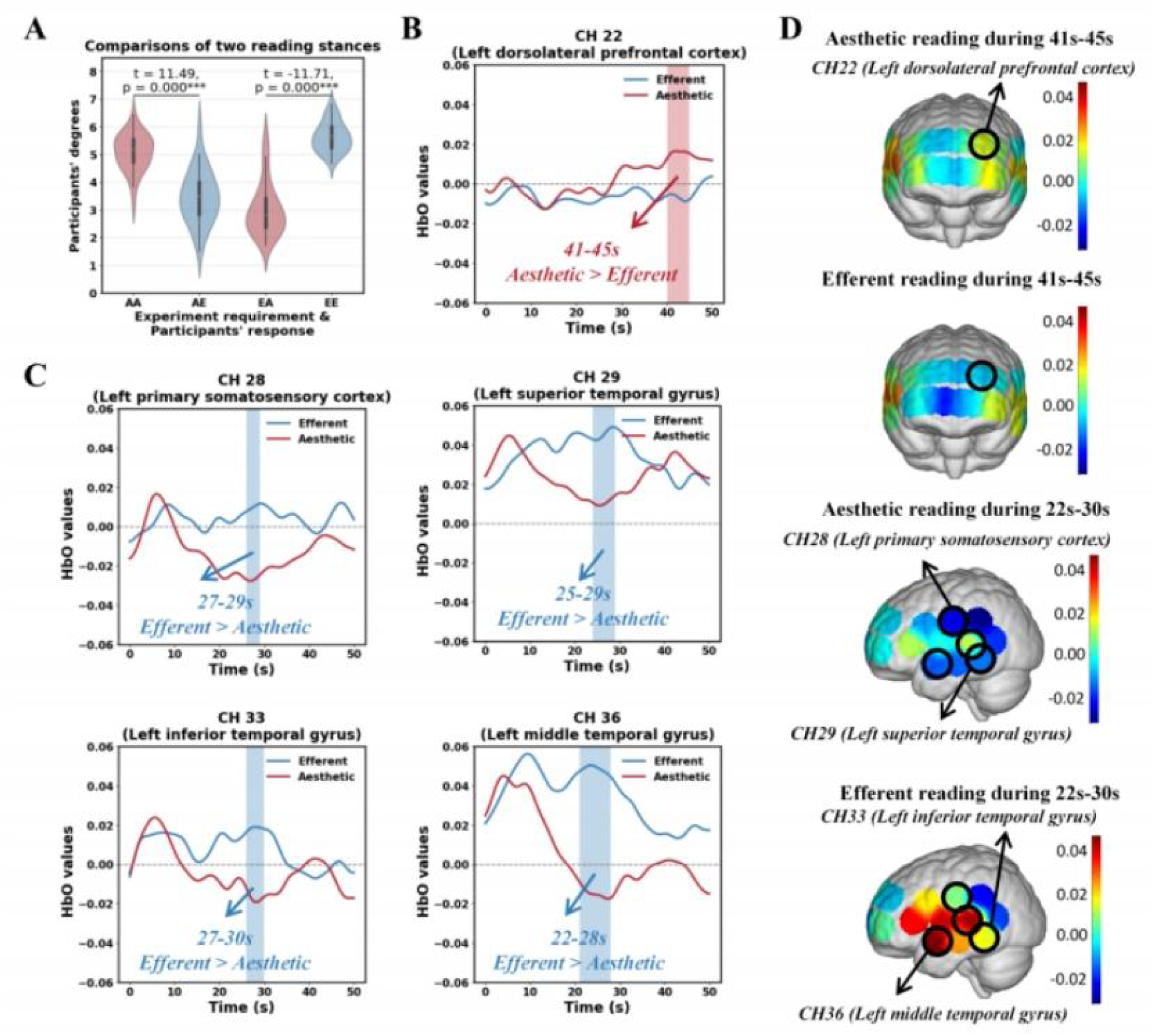
Differences between aesthetic reading and efferent reading. **A,** the distribution of participant’s degree for two reading stances requirements. In the abscissa of the figure, A represents aesthetic reading, and E represents efferent reading. AE indicates the participant’s degree of efferent reading with the experimental requirement of aesthetic reading. **B,** the variation in HbO levels over the 50-second reading period shows significantly higher activation during aesthetic reading compared to efferent reading. At 0s, HbO value was corrected based on the 5-second average value during the resting-state. **C,** the variation in HbO levels over the 50-second reading period shows significantly higher activation during efferent reading compared to aesthetic reading. **D,** Illustrations of the brain regions showing significant differences in activity between aesthetic reading and efferent reading. The black circle highlights areas of statistically significant difference. The red areas indicate higher positive activation, while the blue areas represent greater inhibition.

### 3.2 fNIRS Neuroimaging Results on the Differences between Aesthetic Reading and Efferent Reading

According to Figure 2B and 2C, significant differences in HbO were observed in five channels between aesthetic reading and efferent reading stances: CH22 (left dorsolateral prefrontal cortex), CH28 (left primary somatosensory cortex), CH29 (left superior temporal gyrus), CH33 (left inferior temporal gyrus), and CH36 (left middle temporal gyrus). The detailed information of the t-test results are listed in Appendix A Table A5. From an overall perspective of HbO level changes, aesthetic reading exhibits distinct phased characteristics: based on the real-time trends in HbO concentration, the cognitive process of aesthetic reading can be divided into three stages. During the initial phase (i.e., the first 0-10 seconds), participants exhibited similar brain activity in both aesthetic and efferent reading tasks. CH28, CH29, CH33, and CH36 were simultaneously activated in both reading tasks within the first 0-10 seconds, as determined by the peak and sub-peak HbO levels. Subsequently, in the second phase (i.e., 10-30 seconds), HbO levels in these four channels rapidly decreased during the aesthetic reading task, and were significantly lower than those observed during the efferent reading task (Figure 2C). In detail, CH28 (left primary somatosensory cortex), CH29 (left superior temporal gyrus), CH33 (left inferior temporal gyrus), and CH36 (left middle temporal gyrus) exhibited significantly lower HbO (p<0.05) during aesthetic reading than in efferent reading at the time windows of 27s-29s, 25s-29s, 27s-30s, and 22s-28s, respectively. In the third phase (i.e., 30-50 seconds), HbO activation levels in CH28, CH29, CH33, and CH36 began to rise again during aesthetic reading, eventually reaching levels comparable to those observed during efferent reading in the same phase. Additionally, HbO levels in CH22 (left dorsolateral prefrontal cortex), associated with imaginative processes, was significantly higher during aesthetic reading than efferent reading between 41 and 45 seconds (p < 0.05), the third phase. Specific brain regions involved are illustrated in Figure 2D.

Channels CH28, CH29, CH33, and CH36 display a notable three-phase pattern during aesthetic reading, characterized by an initial increase, followed by a decrease, and then another increase (as illustrated in Figure 3A). These brain regions, associated with semantic comprehension, are likely experiencing a kind of semantic inhibition during the second phase of aesthetic reading. To further test if this semantic inhibition is meaningful to the transcending from language comprehension to aesthetic appreciation, the correlation between each individual’s inhibition level and his/her aesthetic reading status is calculated. Two indexes were first proposed to measure such potential inhibition. The first one was calculated by subtracting the trough value of the second stage from the peak value of the first stage, and was denoted as Δ d1. The second one, as denoted by Δ d2, was derived by subtracting the trough of the second stage from the peak of the third stage (see Figure 3A). The correlations between each Δ d with the corresponding self-report aesthetic levels were shown in Figures 3B and 3C, respectively. The results indicate a positive correlation between Δ d2 in CH28, CH29 and CH33 and aesthetic level (r_CH29_ = 0.485, p < 0.01; r_CH33_ = 0.388, p < 0.05). Therefore, Δ d2, which represents the reactivation level of semantic processing in the third phase after the semantic inhibition in the second phase, may imply higher level of aesthetic appreciation.

**Figure 3.**
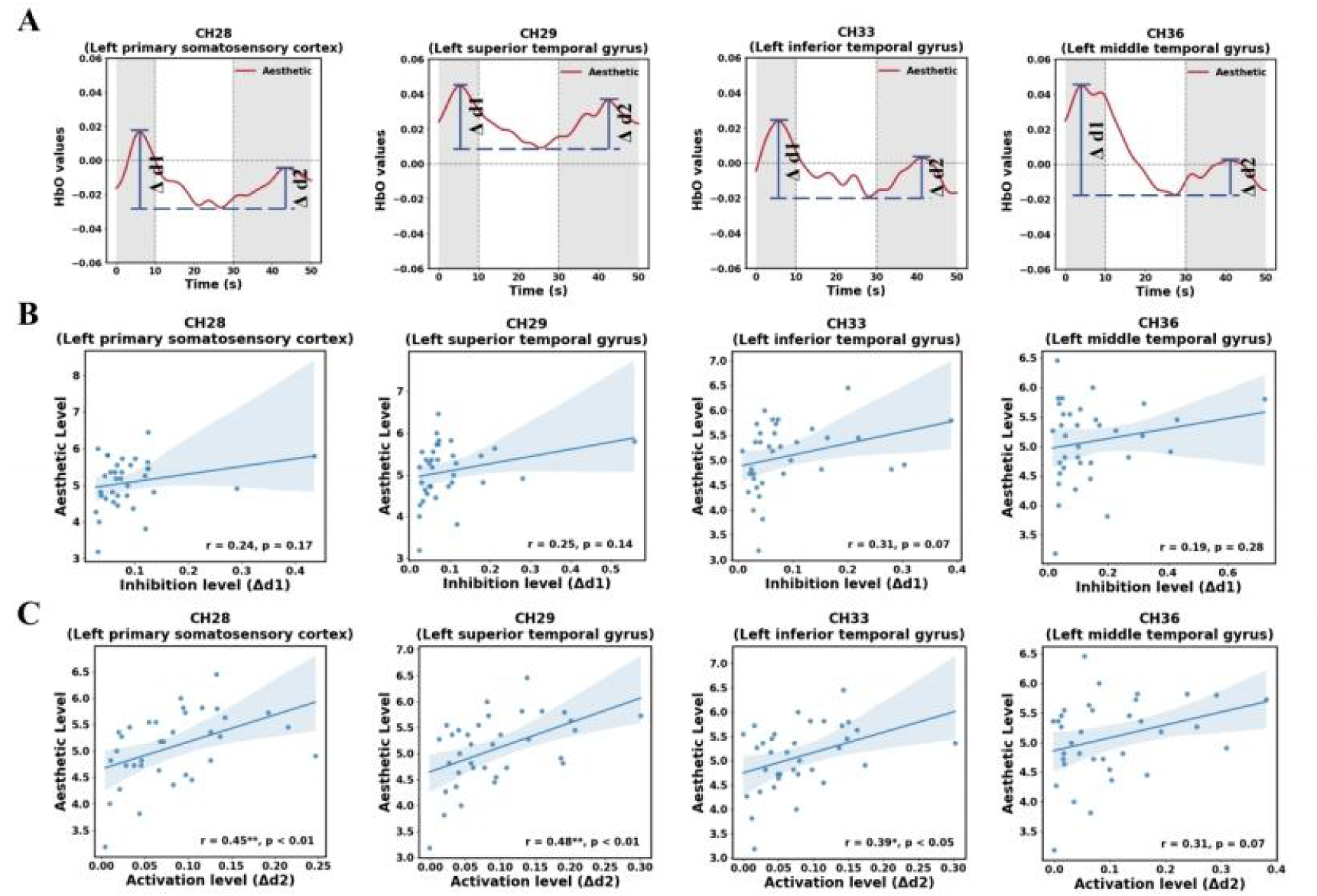
Three-phase changes in HbO during aesthetic reading. **A,** average HbO level changes over the 0-50 second period, with shaded areas representing different stages. **B,** an illustration of the correlation between aesthetic levels and potential semantic inhibition level ( Δ d1). **C,** an illustration of the correlation between aesthetic levels and potential semantic inhibition level ( Δ d2).

## 4 Conclusion and Discussion

Based on aesthetic theories, this study explores the cognitive neural activities involved in the transition from semantic understanding to aesthetic appreciation in Chinese poetry reading, by comparing differences in HbO levels between aesthetic reading and efferent reading across both spatial and temporal dimensions.

Integrating the previous findings in the spatial dimension, this study concludes that aesthetic reading appears to follow a three-phase process. Considering the dynamic changes in the four channels (CH28, CH29, CH33, CH36), we divided the 50-second task into three phases based on the HbO levels during aesthetic reading: an initial rise in HbO levels in the first 10 seconds, a decline between 10 and 30 seconds, followed by a second rise from 30 to 50 seconds. The first phase showed a similar rising trend in HbO levels during both aesthetic and efferent reading. In the second phase, HbO levels in brain regions associated with semantic comprehension decreased during aesthetic reading and were significantly lower than the activation levels observed during efferent reading. In the third phase of aesthetic reading, the activation levels of these four brain regions increase again, ultimately reaching levels comparable to those observed during efferent reading. During this phase, the HBO levels in CH22 (left dorsolateral prefrontal cortex) rise notably, with activation levels significantly exceeding those recorded during efferent reading in the same region.

The CH28 (left primary somatosensory cortex), CH29 (left superior temporal gyrus), CH33 (left middle temporal gyrus), and CH36 (left inferior temporal gyrus) possibly reflect word processing tasks, processing written language, language comprehension, semantic knowledge storage, and semantic control (Aryani, 2018; Binder, 2009; Booth, 2001; Booth, 2006; Gaillard, 2001; Joseph, 2001; Whitney, 2011). The left dorsolateral prefrontal cortex possibly reflects memory (Blumenfeld, 2006; Blumenfeld, 2011; Balconi, 2013), imagination (McNorgan, 2012), empathy (Brothers, 1990; Light, 2009; Bernhardt, 2012; Engen, 2013; Yang, 2018), and aesthetic evaluation (Boccia, 2016). Moreover, the difference in HBO activation levels between the third and second phases in brain regions associated with semantic comprehension (CH28, CH29, CH33) can significantly predict the level of aesthetic appreciation. The brain activities during aesthetic reading are compared with those in efferent reading, summarized in Table 1.

**Table 1.** Brain activities during aesthetic reading, compared with efferent reading.

|  | Phase one<br>(0-10s) | Phase two<br>(10-30s) | Phase three<br>(30-50s) |
| --- | --- | --- | --- |
| Regions associated with semantic understanding<br>(CH28, CH29, CH33, CH36) | Similar | Inhibition | Reactivation |
| Region associated with memory, imagination and empathy<br>(CH22) | Similar | Similar | Activation |

Combining these observations, a three-stage procedure is proposed: semantic understanding, aesthetic contemplation, and aesthetic appreciation. In the semantic understanding phase, readers primarily focus on perceiving and comprehending the content of the text. As they move into the aesthetic contemplation stage, semantic understanding is momentarily suppressed, while aesthetic processing has yet to fully activate. In the final stage of aesthetic appreciation, brain regions associated with memory, imagination, and empathy become active, and the brain areas linked to semantic understanding re-engage. This suggests that, after intense semantic processing, readers adopting an aesthetic stance gradually suppress their attention to non-personalized, abstract semantic information, shifting towards a more personalized and subjective aesthetic experience. The reactivation of semantic understanding in this final phase may be related to memory, imagination, and empathy, rather than the initial comprehension of the text, as this reactivation correlates with self-reported aesthetic appreciation. In fact, imagination and empathy are considered forms of experiential understanding, representing a deeper comprehension of the aesthetic object and grasping its inner meaning (Gadamer, 1989).

The findings of potential semantic inhibition during the transition from external reference to internal response, along with the reactivation of semantic processing intertwined with imagination and empathy, echo several established theories that open the discussion further. For instance, Schiller posits that humans are divided between two fundamental drives: the sensuous drive which is grounded in our physical and emotional nature, and the formal drive, which is associated with reason and rationality. When one of these drives dominates the other, individuals either become overly rational and cold, losing their connection to the natural and emotional world, or become enslaved to their passions, lacking direction and higher purpose. Schiller believes that aesthetic education offers a pathway to achieving a harmonious state of both rationality and sensibility, which is not just a psychological balance but also a moral and cultural ideal. This balance, he argues, fosters both personal and societal freedom (Schiller, 2016). Greene (2000) also proposed that aesthetic education emphasizes the integration of emotional engagement with cognitive processes during reading. This dual engagement can enhance understanding and appreciation of texts. From this perspective, the semantic inhibition and reactivation observed in this study could be viewed as subtle mechanisms for balancing sensibility with rationality, as the latter inherently relies on language.

In Taoism, language is seen as limited in expressing deeper truths, especially the Tao. Zhuangzi states that when ideas are grasped, one should forget the words used to express them. “Those who know the Tao do not speak of it; those who speak of it do not know it, and thus the sage imparts his wisdom without words” (Zhuang, 2015). Similarly, the Chan School asserts that the ultimate truth is beyond both language and rationality, and cannot be fully expressed through words, but can only be realized through enlightenment (Kai, 2017). Language, in this view, acts as a ladder or pathway toward ultimate truth, rather than the final expression of it. Thus, transcending from linguistic comprehension to a higher level of aesthetic experience, involving a deep and harmonious integration of rationality and sensibility, seems to be a common thread in these philosophical perspectives.

This has important implications for further understanding the process of aesthetic reading and, consequently, for improving educational practices. In contemporary poetry teaching, some educators place excessive emphasis on the understanding of word meanings and semantics, focusing on the analysis of syntax and literary technique (Hu, 2010). As a result, students may become detached from their internal experiences and aesthetic engagement, which can negatively impact their aesthetic experience. This study, from a neurophysiological perspective, confirms that in order to genuinely experience aesthetics during reading, it is crucial to undergo a process of semantic inhibition. Only when students disengage from an overly analytical focus on words and syntax can they resonate more deeply with the author, thereby achieving a higher level of understanding and truly appreciating the “beauty” within the text.

It is also important to notice that, although this study has revealed the overall neurophysiological changes across different reading modes, the process of aesthetic reading involves more complex mechanisms, such as memory recall, association, and empathy. Further investigation into the finer neurophysiological processes underlying these mechanisms is needed. Second, this study has preliminarily identified inhibitory phenomena in brain regions associated with semantic processing during aesthetic reading using fNIRS. However, the investigation of brain regions has been limited to the cortical areas. Future research should employ techniques with higher spatial resolution, such as functional magnetic resonance imaging (fMRI), to conduct a more in-depth exploration and further investigate the neurophysiological mechanisms of aesthetic reading.

In summary, this study holds significant interdisciplinary implications. By applying cutting-edge cognitive neuroscience tools and methods, and employing a rigorous randomized controlled trial, the study is the first to comprehensively reveal the temporal and spatial dynamics of brain activity during aesthetic reading. The findings provide new scientific evidence supporting existing theories of aesthetic reading— particularly those concerning the nature of aesthetic experience and the two-stage model of text comprehension and aesthetic experience—while enhancing the credibility of these theories. More importantly, the study makes a substantial contribution by uncovering novel empirical evidence that addresses gaps in previous theories, offering valuable insights for theoretical innovation. Thus, this study calls for further interdisciplinary research, particularly encouraging the use of novel data and methodologies to foster interdisciplinary dialogue and advance our understanding of educational principles.

## Ethics Statement

The studies involving human participants were reviewed and approved by the local ethics committee at the School of Social Sciences, Tsinghua University.

## Declaration of Interests

The authors declare that they have no known competing financial interests or personal relationships that could have appeared to influence the work reported in this paper.

## Data and materials availability

All data are available from the corresponding author upon reasonable request.

## Acknowledgment

This work was supported by the National Natural Science Foundation of China (62177030), the Beijing Educational Science Foundation of the Fourteenth 5-year Planning (BGEA23019), the Tsinghua University Initiative Scientific Research Program (20225080045), the Education Innovation Grants of Tsinghua University (DX05_02), and the Teaching Reform Project of the Instruction Committee of Psychology in Higher Education by the Ministry of Education of China (20222008).

## Notes

### Competing Interest Statement

The authors have declared no competing interest.

### Summary of Updates

The introduction, results, and discussion sections of the article have been updated, with the addition of Figure 3 to supplement the verification results.

