## Supplemental Information for "The Role of Semantic Inhibition in Aesthetic Appreciation: fNIRS Evidence from Poetry Reading"

**Appendix A: Tables**

| **Table A1: Initial Ratings (1-7 Scale) for the 25 Preselected Poems** | | | | | | | |
| --- | --- | --- | --- | --- | --- | --- | --- |
| **Poem title** | **Difficulty** | **Familiarity** | **Emotional Valence** | **Level of Engagement** | **Aesthetic Response** | **Imaginative Difficulty** | **Selected for experiment** |
| 题破山寺后禅院 | 2.62 | 5.46 | 4.46 | 3.69 | 4.85 | 2.46 | √ |
| 春夜喜雨 | 1.46 | 6.69 | 5.38 | 4.54 | 4.77 | 2.77 | √ |
| 辋川闲居赠裴秀才迪 | 2.77 | 2.54 | 4.38 | 3.77 | 4.69 | 2.85 | √ |
| 湖口望庐山瀑布泉 | 2.62 | 1.92 | 4.69 | 3.92 | 4.62 | 2.54 | √ |
| 泛永嘉江日暮回舟 | 3.69 | 1.38 | 4.23 | 2.77 | 4.54 | 3.00 | √ |
| 旅夜书怀 | 2.15 | 5.38 | 2.31 | 3.85 | 4.54 | 2.38 | √ |
| 渡荆门送别 | 1.77 | 5.92 | 3.54 | 3.54 | 4.31 | 2.31 | √ |
| 终南别业 | 2.15 | 4.85 | 4.85 | 3.77 | 4.23 | 2.69 | √ |
| 野望 | 2.92 | 3.38 | 3.00 | 3.46 | 4.23 | 3.15 | √ |
| 旅宿 | 2.54 | 1.85 | 2.08 | 3.54 | 4.15 | 2.85 | √ |
| 水槛遣心 | 2.77 | 2.54 | 4.62 | 3.23 | 4.00 | 2.54 | √ |
| 耶溪泛舟 | 2.23 | 3.23 | 3.85 | 2.92 | 3.77 | 2.77 | √ |
| 月夜忆舍弟 | 2.23 | 5.08 | 2.08 | 3.46 | 3.77 | 2.46 | √ |
| 终南山 | 3.46 | 2.38 | 4.15 | 3.23 | 3.69 | 3.38 | √ |
| 和晋陵陆丞早春游望 | 2.85 | 2.54 | 2.54 | 3.31 | 3.69 | 2.77 | √ |
| 登岳阳楼 | 2.62 | 4.46 | 2.31 | 4.00 | 3.38 | 2.77 | √ |
| 从军行 | 2.69 | 4.46 | 4.08 | 4.54 | 3.31 | 3.00 | √ |
| 留别王侍御维 | 2.54 | 2.69 | 2.31 | 2.92 | 3.23 | 3.00 | √ |
| 新年作 | 2.38 | 2.08 | 2.23 | 2.92 | 3.15 | 3.00 | √ |
| 夜泊牛渚怀古 | 2.54 | 2.46 | 2.62 | 2.85 | 3.08 | 3.38 | √ |
| The average scores for the 20 experimental materials | 2.55 | 3.56 | 3.49 | 3.51 | 4.00 | 2.80 |  |
| 晚泊浔阳望庐山 | 2.77 | 1.85 | 3.69 | 2.38 | 3.08 | 3.38 |  |
| 与诸子登岘山 | 3.15 | 2.85 | 2.31 | 3.31 | 3.00 | 3.23 |  |
| 塞下曲·其四 | 3.15 | 2.00 | 2.08 | 3.08 | 2.69 | 3.46 |  |
| 喜见外弟又言别 | 2.38 | 2.46 | 3.00 | 4.15 | 2.69 | 2.85 |  |
| 蜀先主庙 | 2.77 | 2.15 | 2.69 | 3.00 | 2.62 | 3.38 |  |
| The average scores for the 5 unselected materials | 2.84 | 2.26 | 2.75 | 3.18 | 2.82 | 3.26 |  |

| **Table A2: Level of Engagement and Emotional Valence (N=35)** | | | | |
| --- | --- | --- | --- | --- |
|  | Minimum | Maximum | Mean | Standard Deviation |
| Level of Engagement | 2 | 7 | 4.81 | 1.11 |
| Emotional Valence | 3 | 6 | 4.47 | 0.87 |

| **Table A3: Comparison of Familiarity Levels among Four Poem Sets** | | | | |
| --- | --- | --- | --- | --- |
| Group | Mean | Standard Deviation | F | *p* |
| Group A1 | 3.63 | 2.565 | 0.09 | 0.96 |
| Group A2 | 3.63 | 2.362 |  |  |
| Group B1 | 4.00 | 2.449 |  |  |
| Group B2 | 3.63 | 2.553 |  |  |

| **Table A4: T-test for Differences in Multiple-Choice Question Accuracy Rates during Different Reading Tasks (N=70)** | | | | | | |
| --- | --- | --- | --- | --- | --- | --- |
| Group | N | Mean | Standard Deviation | Standard Error of Mean | t | *p* |
| Aesthetic Reading | 35 | 6.29 | 1.69 | 0.29 | -0.30 | 0.76 |
| Efferent Reading | 35 | 6.40 | 1.48 | 0.25 |  |  |

| **Table A5: T-test for Channel Activation Differences Between Aesthetic Reading and Efferent Reading** | | | | | | |
| --- | --- | --- | --- | --- | --- | --- |
| Channel | **Time(s)** | **Aesthetic Reading** | **Efferent Reading** | **Aesthetic Reading - Efferent Reading** | t | ***p*** |
| 22 | 41 | 0.014 | -0.005 | 0.020 | -2.046 | 0.049 |
|  | 42 | 0.016 | -0.005 | 0.022 | -2.095 | 0.044 |
|  | 43 | 0.016 | -0.007 | 0.023 | -2.130 | 0.040 |
|  | 44 | 0.016 | -0.008 | 0.025 | -2.188 | 0.036 |
|  | 45 | 0.016 | -0.009 | 0.024 | -2.168 | 0.037 |
| 28 | 26 | -0.026 | 0.007 | -0.033 | 2.071 | 0.046 |
|  | 27 | -0.027 | 0.008 | -0.036 | 2.261 | 0.030 |
|  | 28 | -0.027 | 0.010 | -0.037 | 2.303 | 0.027 |
|  | 29 | -0.026 | 0.011 | -0.037 | 2.192 | 0.035 |
| 29 | 25 | 0.010 | 0.043 | -0.033 | 2.108 | 0.042 |
|  | 26 | 0.009 | 0.043 | -0.034 | 2.216 | 0.033 |
|  | 27 | 0.010 | 0.045 | -0.036 | 2.316 | 0.027 |
|  | 28 | 0.011 | 0.048 | -0.037 | 2.311 | 0.027 |
|  | 29 | 0.013 | 0.049 | -0.036 | 2.123 | 0.041 |
| 33 | 27 | -0.015 | 0.018 | -0.032 | 2.140 | 0.040 |
|  | 28 | -0.018 | 0.019 | -0.037 | 2.504 | 0.017 |
|  | 29 | -0.019 | 0.019 | -0.038 | 2.512 | 0.017 |
|  | 30 | -0.017 | 0.018 | -0.036 | 2.241 | 0.032 |
| 36 | 22 | -0.011 | 0.048 | -0.059 | 2.133 | 0.040 |
|  | 23 | -0.013 | 0.050 | -0.063 | 2.219 | 0.033 |
|  | 24 | -0.015 | 0.050 | -0.065 | 2.235 | 0.032 |
|  | 25 | -0.015 | 0.050 | -0.065 | 2.186 | 0.036 |
|  | 26 | -0.016 | 0.049 | -0.064 | 2.132 | 0.040 |
|  | 27 | -0.017 | 0.047 | -0.064 | 2.102 | 0.043 |
|  | 28 | -0.017 | 0.046 | -0.063 | 2.050 | 0.048 |

Note: The activation level for aesthetic reading is calculated as the HbO during the aesthetic reading task minus the HbO during the resting stance. Similarly, the activation level for efferent reading is calculated as the HbO during the efferent reading task minus the HbO during the resting stance.

**Appendix B: Supplementary Materials**

**Table B1. Reading Stance Scale**

Please describe your feelings while reading the ten poems just now. Rate your response on a scale of 1 to 7, where 1 means "Strongly Disagree" and 7 means "Strongly Agree."

| **Items** | **1** | **2** | **3** | **4** | **5** | **6** | **7** |
| --- | --- | --- | --- | --- | --- | --- | --- |
| 1. When I read poetry, I focus on understanding the literal meaning of the lines or a particular word. | ○ | ○ | ○ | ○ | ○ | ○ | ○ |
| 2. When I read poetry, I pay attention to the writing techniques of the poem. | ○ | ○ | ○ | ○ | ○ | ○ | ○ |
| 3. When I read poetry, I look for detailed information from the poem. | ○ | ○ | ○ | ○ | ○ | ○ | ○ |
| 4. When I read poetry, I focus on analyzing the genre and style of the poem. | ○ | ○ | ○ | ○ | ○ | ○ | ○ |
| 5. When I read poetry, I focus on understanding the background knowledge of the poem. | ○ | ○ | ○ | ○ | ○ | ○ | ○ |
| 6. When I read poetry, I focus on analyzing the main themes of the poem. | ○ | ○ | ○ | ○ | ○ | ○ | ○ |
| 7. When I read poetry, I feel that I have had similar experiences. | ○ | ○ | ○ | ○ | ○ | ○ | ○ |
| 8. When I read poetry, I can recall the emotions from similar experiences. | ○ | ○ | ○ | ○ | ○ | ○ | ○ |
| 9. When I read poetry, I am reminded of specific details from similar experiences. | ○ | ○ | ○ | ○ | ○ | ○ | ○ |
| 10. While reading, I predict the actions or emotional changes of the characters in the poem. | ○ | ○ | ○ | ○ | ○ | ○ | ○ |
| 11. When reading poetry, I imagine the scenes depicted in the ancient poems. | ○ | ○ | ○ | ○ | ○ | ○ | ○ |
| 12. When I read poetry, I can feel the rich and full emotions of the poet or protagonist. | ○ | ○ | ○ | ○ | ○ | ○ | ○ |
| 13. I am able to empathize with the poet emotionally. | ○ | ○ | ○ | ○ | ○ | ○ | ○ |
| 14. I can understand the actions of the poet or protagonist, creating a sense of identification. | ○ | ○ | ○ | ○ | ○ | ○ | ○ |
| 15. I can stand in the shoes of the poet or protagonist to understand the poem. | ○ | ○ | ○ | ○ | ○ | ○ | ○ |
| 16. After reading, I am able to find answers to some practical life problems. | ○ | ○ | ○ | ○ | ○ | ○ | ○ |
| 17. When I read poetry, I make aesthetic evaluations of the content of the text. | ○ | ○ | ○ | ○ | ○ | ○ | ○ |
